# Genetic dissection of the tissue-specific roles of type III effectors and phytotoxins in the pathogenicity of *Pseudomonas syringae* pv. *syringae* to cherry

**DOI:** 10.1101/2024.02.06.578989

**Authors:** Andrea Vadillo-Dieguez, Ziyue Zeng, John W. Mansfield, Nastasiya F. Grinberg, Samantha C. Lynn, Adam Gregg, John Connell, Richard J. Harrison, Robert W. Jackson, Michelle T. Hulin

**Affiliations:** NIAB, Lawrence Weaver Road, Cambridge, CB3 0LE, UK; Imperial College London, London, SW7 2BX, UK; School of Biosciences and the Birmingham Institute of Forest Research, University of Birmingham, Birmingham, B15 2TT, UK

**Keywords:** *Pseudomonas syringae*, type 3 effectors, phytotoxins, mutagenesis, virulence, *Prunus*, comparative genomics

## Abstract

When compared with other phylogroups (PGs) of the *Pseudomonas syringae (Ps)* species complex, *Ps* pv. s*yringae* strains within PG2 have a reduced repertoire of type III effectors (T3Es) but produce several phytotoxins. Effectors within the cherry pathogen *Pss*9644 were grouped based on their frequency in strains from *Prunus* as: the conserved effector locus (CEL) common to most *Ps* pathogens; a CORE of effectors common to PG2; a set of PRUNUS effectors common to cherry pathogens; and a FLEXIBLE set of T3Es. *Pss*9644 also contains gene clusters for biosynthesis of toxins syringomycin/syringopeptin and syringolin A. After confirmation of virulence gene expression, mutants with a sequential series of T3E and toxin deletions were pathogenicity tested on wood, leaves and fruits of sweet cherry (*Prunus avium*) and leaves of ornamental cherry (*Prunus incisa*). The toxins had a key role in disease development in fruits but were less important in leaves and wood. An effectorless mutant retained some pathogenicity to fruit but not wood or leaves. Striking redundancy was observed amongst effector groups. The CEL effectors have important roles during the early-stages of leaf infection and acted synergistically with toxins in all tissues. Deletion of separate groups of T3Es had much more effect in *Prunus incisa* than in sweet cherry. Mixed inocula were used to complement the toxin mutations *in trans* and indicated that strain mixtures may be important in the field. Our results highlight the niche-specific role of toxins in cherry tissues and the complexity of effector redundancy in the pathogen *Pss*9644.

## Introduction

Bacterial pathogenicity to plants has, for many diseases, been closely linked to the secretion of effector proteins (Lovelace et al., 2023). Genes encoding effector proteins that are injected into host cells by the Type III secretion system (T3SS) were originally cloned from plant pathogenic bacteria, not by their virulence function, but by their ability to act as avirulence (*avr*) genes whose products triggered the hypersensitive resistance reaction (effector triggered immunity, ETI; Jones and Dangl, 2006, Lovelace et al., 2023, Mansfield, 2009, Xin et al., 2018). The initial isolation of *avr* genes was based on the exchange of genomic libraries between races of pathogens that displayed differential virulence on certain varieties of crop plants such as soybean, pea and pepper (Mansfield, 2009, Staskawicz et al., 1984). The *avr* genes identified were often absent from virulent races of the pathogens and therefore were not at first assigned essential roles in basic pathogenicity. Effectors (T3Es), produced by plant pathogens are now recognised to have key roles in the suppression of host defences, both ETI and those triggered by microbe-associated molecular patterns (MAMPs, MTI). They also have roles in the creation of conditions *in planta* that benefit microbial colonisation (Ekanayake et al., 2022, Lovelace and Ma, 2022, Lovelace et al., 2023, Nomura et al., 2023, Xin et al., 2018) . Some effectors have been predicted to have enzymatic activity that is required for virulence functions (Grant et al., 2006, Washington et al., 2016).

There are now several examples of T3Es that have individually been identified as being required for pathogenicity in bacterial plant pathogens, for example VirPphA (also known as HopAB1) in *Pseudomonas syringae* pv. *phaseolicola* (Pph, Jackson et al., 1999), DspA/E in *Erwinia amylovora* (Bogdanove et al., 1998, Yuan et al., 2021) and AvrE, AvrPtoB and AvrPto in strains of *Ps.* pv. *tomato* (*Pto, Xin et al., 2018).* However, deletion of a single effector more often fails to reduce disease following artificial inoculation. The presence of redundant effector groups (REGs) was clarified by the landmark studies on *Pto* strain DC3000 by Collmer and colleagues (Cunnac et al., 2011, Kvitko et al., 2009, Wei and Collmer, 2018). Deletion of individual effectors from REGs did not cause a loss of pathogenicity, leading to the description of effectors as “*collectively essential but individually dispensable”* (Kvitko et al., 2009).

The conserved effector locus (CEL), containing two to four effector genes *hopAA1*, *avrE1*, *hopM1* and *hopN1,* has emerged as an important common determinant of pathogenicity to leaves in several strains of *Pto* and *Ps*. pv. *actinidiae (Psa)* (Alfano et al., 2000, Dillon et al., 2019, Jayaraman et al., 2020). Certain effectors have been assigned functions for promotion of symptom formation rather than the promotion of initial bacterial colonisation, for example HopAM1-1, HopG1 and HopM1 in DC3000 (Badel et al., 2006, Cunnac et al., 2011). In addition to effector proteins, many strains of *Ps* also secrete a second class of pathogenicity factors, low molecular weight phytotoxins such as coronatine, phaseolotoxin and syringomycin that also have key roles in symptom production but are not always required for bacterial multiplication *in planta* (Bender et al., 1999, Geng et al., 2014, Scholz-Schroeder et al., 2001).

*Pseudomonas syringae* is an example of a species complex within which pathogenicity to certain host plants has been linked through bioinformatic analyses to the presence of specific T3E repertoires (Baltrus et al., 2017, Newberry et al., 2019). A good example of this is the economically important bacterial canker disease of *Prunus*, which is caused by members of at least six different clades of *Ps*. The main causal agents of cherry canker are *Ps*. pv. *morsprunorum* (*Psm*) races 1 (*Psm1*) and 2 (*Psm2*) and *Ps.* pv. *syringae* (*Pss*). Despite their differential core genomes, comparative genomics using Bayes-Traits analysis, has identified convergent patterns of gain and loss of effectors associated with clades of *Ps* causing canker, notably the gain of *hopAR1, hopBB1, hopBF1,* and *hopH1* (Hulin et al., 2018a). Strains of *Psm1* and *Psm2* encode numerous effector proteins (from 30 -35). By contrast, strains of *Pss* have fewer effectors (15-18), but, unlike *Psm*, encode biosynthetic clusters for up to four phytotoxins: syringomycin, syringopeptin, syringolin A and mangotoxin. The role of syringomycin and syringopeptin in the production of necrotic lesions on cherry fruits has been analysed by Scholz-Schroeder et al. (Scholz-Schroeder et al., 2001). It has been suggested that the production of phytotoxic metabolites might compensate for the low numbers of effectors in *Pss* (Hulin et al., 2020, Xin et al., 2018). Indeed, the importance of these toxins was emphasised in a recent genome-wide mutagenesis screen by Helmann et al., 2019 on bean. They identified syringomycin as one of the most important fitness determinants of the related bean pathogen *Pss* B278A in the bean apoplast.

Symptoms of bacterial disease of *Prunus* are observed on leaves, buds, fruits and woody tissues (Hulin et al., 2018b, Hulin et al., 2020). Recent screening experiments have identified resistance to the canker pathogens in leaves of wild cherry and related ornamental *Prunus* species (Hulin et al., 2022, Lienqueo et al., 2024). Resistance to *Pss* in *Prunus incisa* was found to be dosage dependent, being overcome by infiltration with high inoculum concentrations (> 10^8^ per ml of infiltrated suspension).

We carried out a genetic dissection of the role of effectors and toxins to understand the ability of the phylogroup 2 strain *Pss*9644 to invade and cause symptoms in woody shoots, fruits and leaves, of sweet cherry, *Prunus avium.* The roles of the pathogenicity factors were also assessed in leaves of *Prunus incisa.* Following the identification of groups of effector genes, including some that were physically unlinked in the genome but common to pathogens of *Prunus*, successive rounds of deletion mutagenesis led to the construction of effectorless and toxinless mutants. Creation of the panel of mutants has allowed the association between the presence of certain T3Es and pathogenicity to be examined, and the following hypotheses to be tested:

1. Redundant effector groups exist in *Pss*9644.
2. Toxins and effectors act synergistically to promote infection.
3. Toxins and effectors vary in their impact on pathogenicity in different cherry tissues.
4. The effectors required to cause symptoms in *Prunus incisa* differ from those essential for virulence to *Prunus avium*.

## Results

### Categorisation of effectors and toxins in phylogroup 2

*P. syringae* phylogroup 2 is a diverse clade within the *Ps* species complex that contains pathogens of most major crop species. A Maximum Likelihood phylogeny based on the core genome placed 74 strains isolated from sweet cherry across the phylogeny in three clades (PG2a, PG2b and PG2d). The strain used in this study, *Pss*9644, is an isolate from cherry within phylogroup 2d (PG2d), previously characterised to be pathogenic to woody tissues, fruit and leaves (Hulin et al., 2018b). We re-sequenced its complete genome revealing one chromosome (6,164,862 bp) and a small plasmid (45,481 bp). Putative T3Es present in *Pss*9644 and a wider range of cherry pathogen genomes were identified by homology and classified into four different categories according to their frequency in cherry pathogen genomes (Figure 1). The conserved effector locus (CEL) is found in most strains of pathogenic *Ps,* and in *Pss*9644 comprised *hopAA1, hopM1,* and *avrE1*. A second group designated CORE (C) effectors was common to other PG2 strains, *hopAG1, hopAH1, hopAI1, and hopI1*. Thirdly, PRUNUS (P) effectors were commonly found in strains isolated from cherry and other *Prunus* spp., *hopAR1, hopH1, hopA2, hopW, hopAW1 and avrRpm1*. Finally, a group defined as FLEXIBLE (F) was variously distributed amongst PG2 strains, *hopAF1, hopAZ1, and hopBE1*. In addition, *Pss*9644 was found to contain gene clusters for the biosynthesis of the toxins syringomycin, syringopeptin (*syrsyp*), and syringolin A (*sylA*).

**Figure 1.**
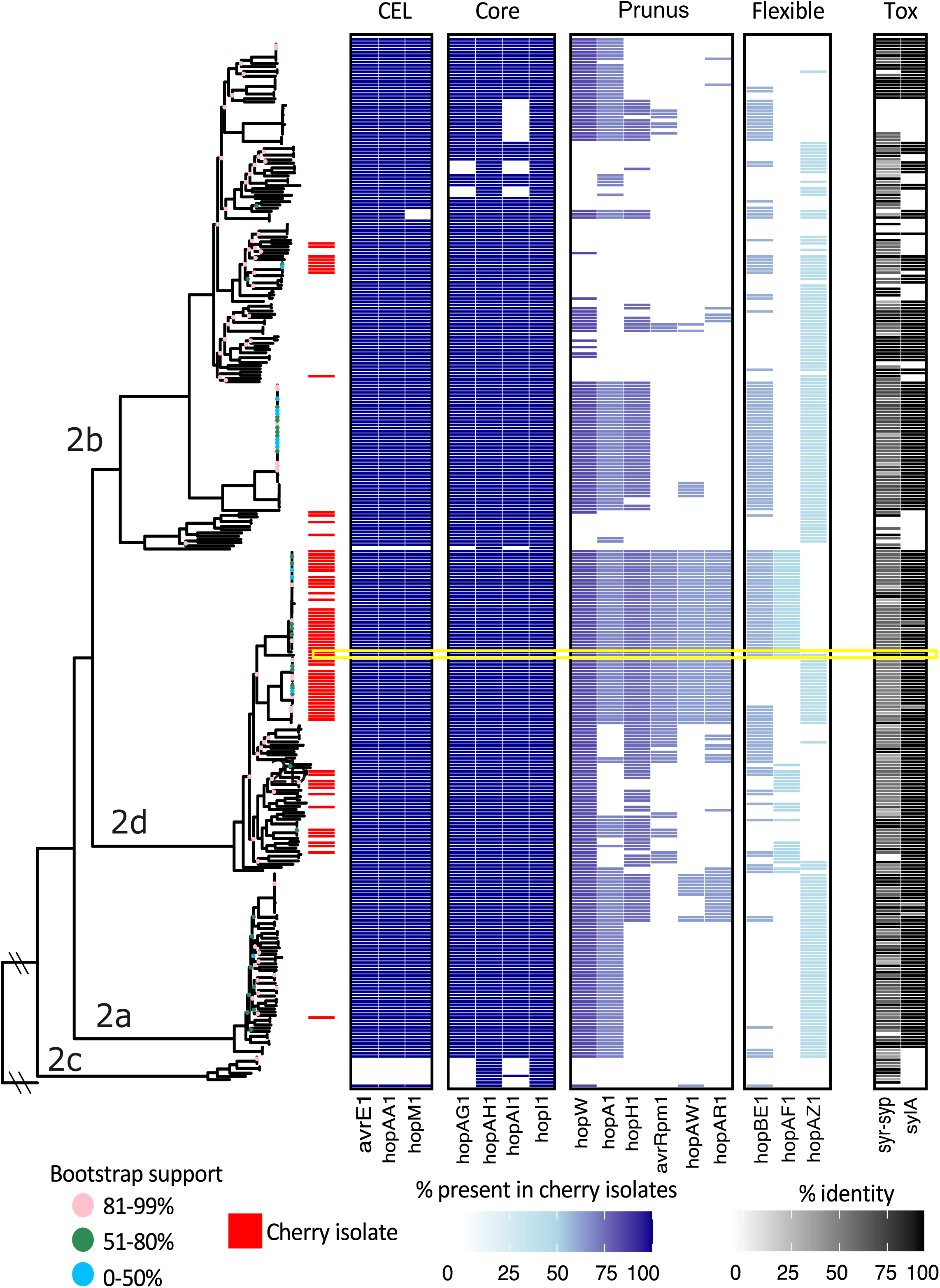
Maximum Likelihood phylogeny of *Ps* strains in phylogroup 2 based on the core genome according to bootstrap support 0-50%; 51-80% and 81-99%. Four different categories of T3E are shown in shades of blue according to their frequency in genomes of isolates from cherry (red): CEL, CORE, PRUNUS, FLEXIBLE. Presence of *syrsyp* and *sylA* clusters are represented with shades of black according to the % identity to the reference in antiSMASH. *Pss*9644 is highlighted with a yellow rectangle. *Pph*1448A was used as an outgroup.

### T3E and phytotoxin genes are upregulated in *hrp*-inducing medium

The expression of genes in *Pss*9644 identified to encode T3Es and enzymes predicted to be involved in toxin synthesis was examined using RNA sequencing (Figure 2). Gene expression was observed in *Pss*9644 grown to exponential phase in rich medium (Kings medium B, KB) and in *hrp* (hypersensitive resistance and pathogenicity)-inducing minimal medium (HMM), which is a simple mimic of the *in planta* environment. The transcripts detected by RNAseq are reported in Supplementary Figure S1. All the predicted genes for T3Es and toxin biosynthesis were expressed in both media. The upregulated expression in HMM compared with KB varied significantly between one to six-log2-fold change except for *hopAI1*. For example, *hopAR1*, *hopAA1* and *avrE1* were very strongly induced in the minimal medium. The weakest relative expression was identified for *hopAF1*, *hopAW1 and hopW* with around one-log2-fold change. In addition, *hopAI1*, in the operon *hopAG1-hopAH1-hopAI1* in CORE effector group, was not differentially expressed. Genes involved in toxin synthesis were also expressed more strongly in HMM medium.

**Figure 2.**
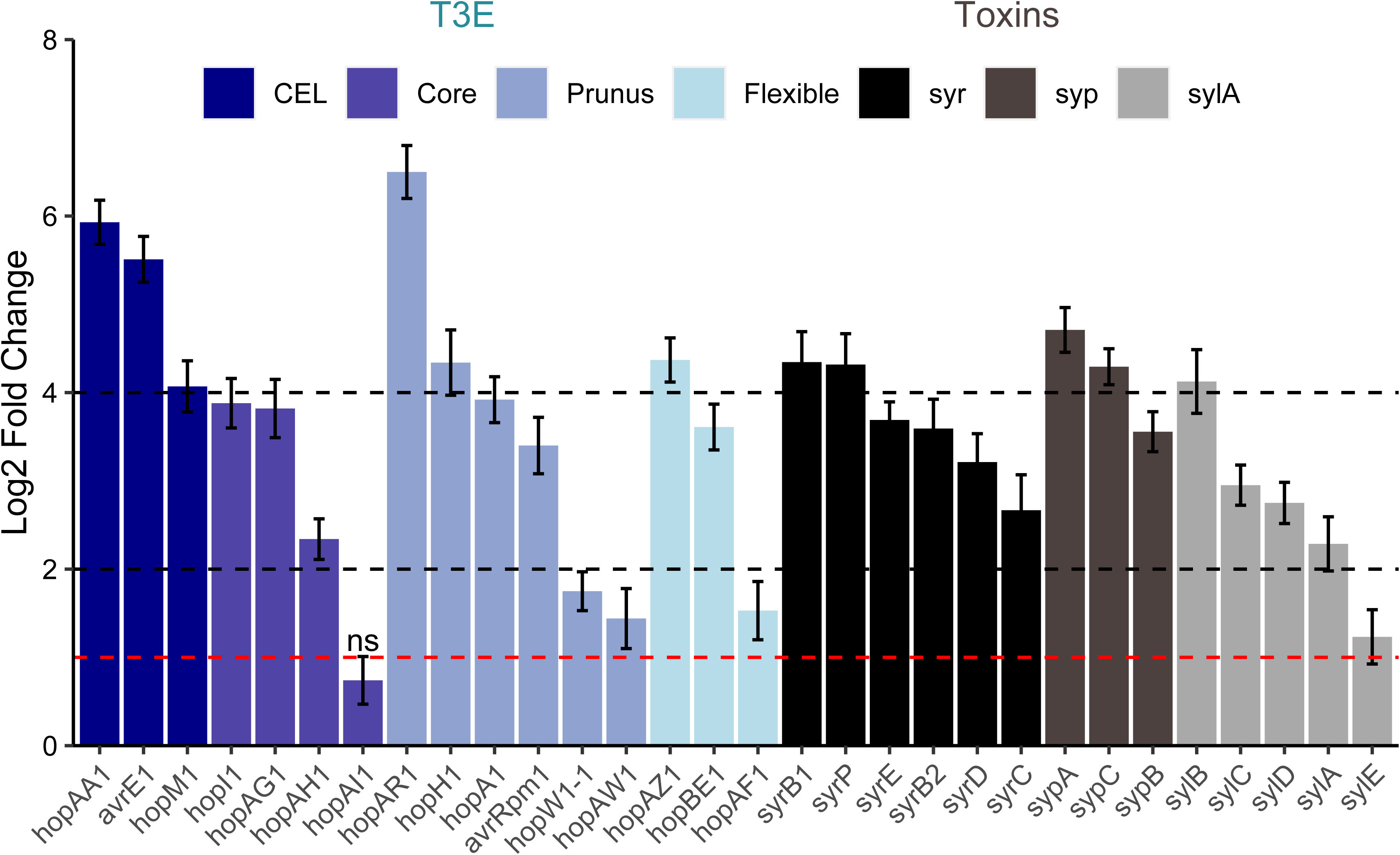
Log 2-Fold change ratio of the upregulated expression of genes encoding effectors and toxin synthesis in *hrp*-inducing minimal medium (HMM) compared to King’s medium B (KB) in *Pss*9644 wild-type strain. T3Es in the four categories of Fig. 1 are colour coded in shades of blue according to their frequency in phylogroup 2 and genomes of isolates from cherry: CEL, CORE, PRUNUS andf FLEXIBLE. Toxin clusters are highlighted in shades of black. Legend represents *syr*, syringomycin; *syp*, syringopeptin; *sylA*, syringolin A. Lines represent the Log 2-Fold change threshold of 1, 2 and 4. ns: non-significant (log2fc < 1). Values represent the average of three biological replicates and error bars represent the standard error of the mean; the experiment was performed once.

### Pathogenicity tests identify tissue-specific effects for virulence factors

Given that most effectors and toxins were differentially expressed in HMM, we predicted that these genes were likely to play an important role in the pathogen’s ability to cause disease in cherry. Deletion mutants as listed and named in Table 1, were constructed sequentially according to the T3E group frequency in cherry isolates from FLEXIBLE to CEL. They were then compared with wildtype *Pss*9644 to determine the effects of mutations on pathogenicity in sweet cherry woody tissues, fruits, and leaves, and in leaves of *Prunus incisa*.

**Table 1.**
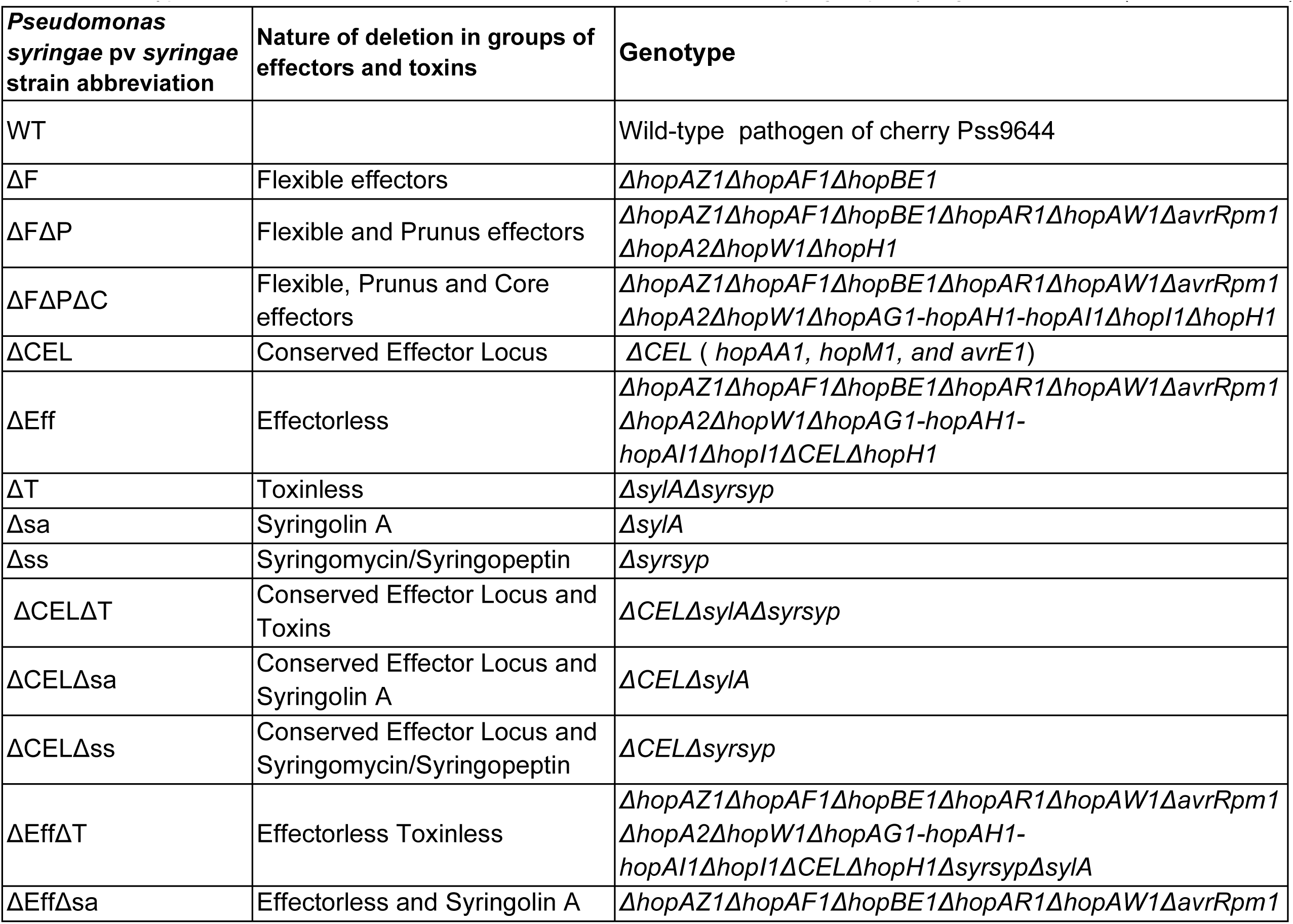
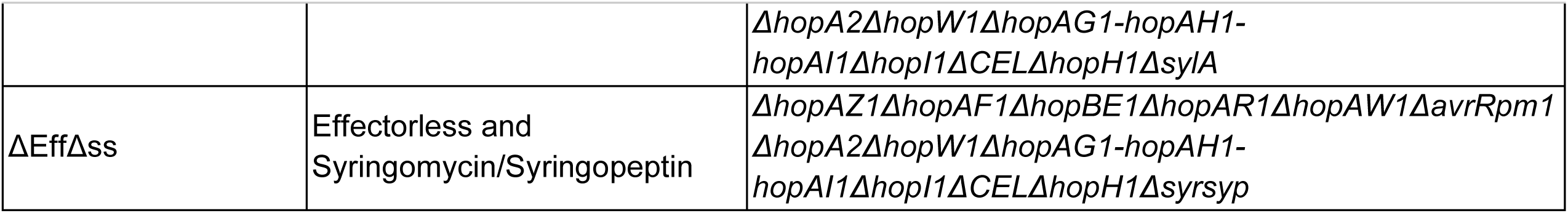
Genotypes of the effector and toxin deletion mutants created in *P. syringae* pv. *syringae* strain 9644 (Roberts, 2012).

### Most effectors, but not toxins, are required to cause disease symptoms in wood

Pathogenicity was compared in both cut shoot and whole tree inoculation assays (Fig. 3). In woody stems results were more variable than in other tissues. In cut shoot assays (Fig.3a), the wildtype *Pss*9644 caused an average lesion length of 9 mm. No significant differences in lesion size was observed when shoots were inoculated with the ΔCEL, Δss, Δsa, ΔT, ΔF or ΔFΔP mutants. However, the ΔFΔPΔC triple mutant caused a lesion significantly smaller than the wildtype. Surprisingly, the CEL deletion mutant (ΔCEL) caused lesions very similar to wildtype. The deletion of both toxins (ΔT) also had a minor effect but the combination of CEL and Toxin deletions in (ΔCELΔT) greatly reduced lesion lengths. The deletion of all effector clusters i.e. ΔCELΔFΔPΔC, created an effectorless (ΔEff) mutant that was the least pathogenic strain tested.

**Figure 3.**
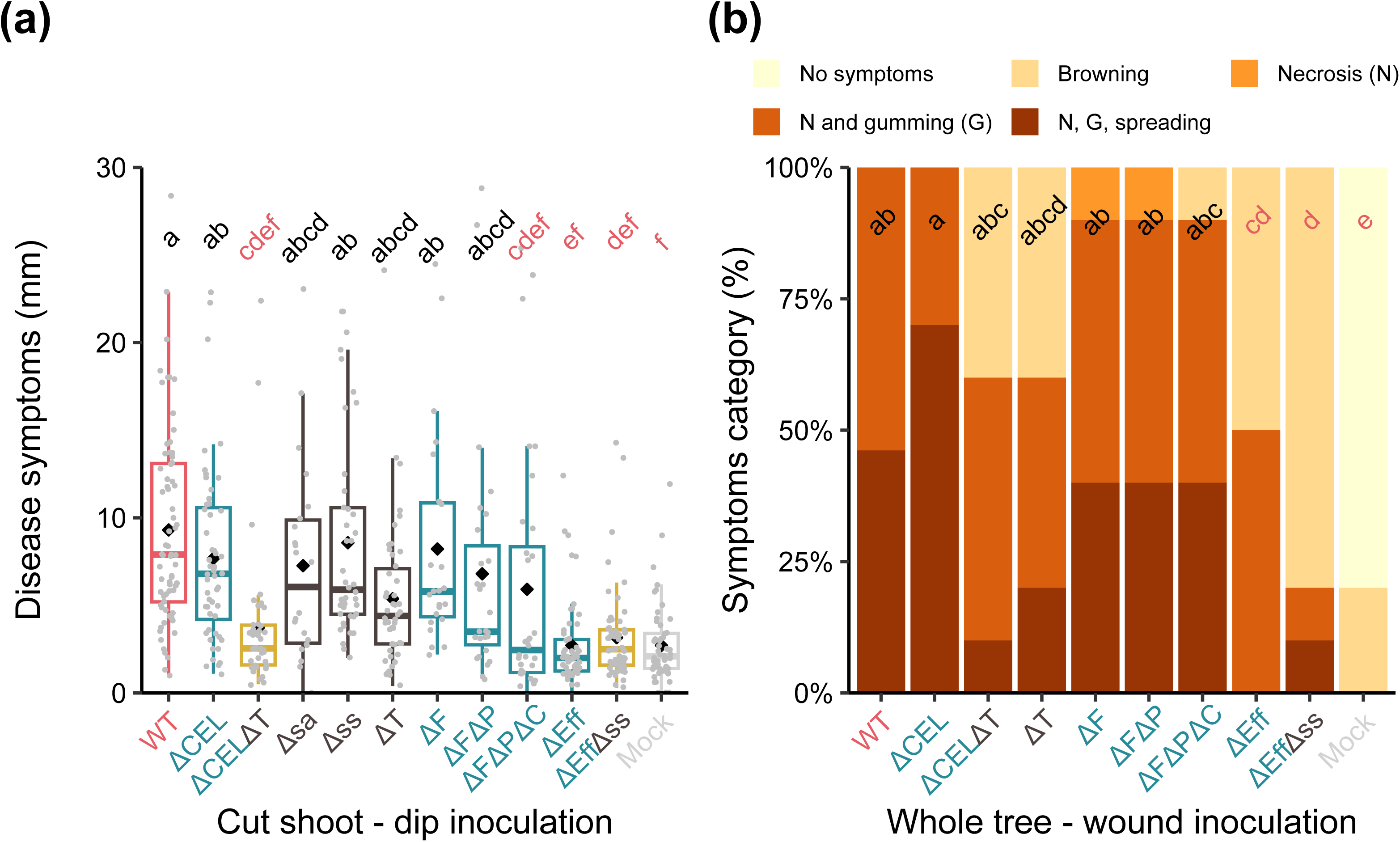
Lesion formation in cut shoots and woody stems on whole trees of cv. Sweetheart inoculated with wild-type *Pss*9644 and deletion mutants. Mutants are described in detail in Table 1. WT: wild-type, CEL: Conserved Effector Locus, F: FLEXIBLE group, P: PRUNUS group, C: CORE group, Eff: all effectors in CEL, FLEXIBLE, PRUNUS, CORE groups, sa: syringolin A cluster, ss: syringomycin/syringopeptin cluster, T: both toxin clusters. Mock:10 mM MgCl_2_. Letters in common above data points indicate no significant difference between treatments. Letters in red indicate significant differences compared to wildtype (p<0.05). Representative symptoms are shown in Fig. S3. (a) Length of lesions produced in cut shoots. Data from three repeated experiments with 45 shoots in total for each treatment were analysed after log transformation by ANOVA and posthoc Tukey-Kramer HSD tests to assess pairwise differences between mutants. (b) Percentage of inoculations (n=10) in each disease score category after wound inoculation into trees. Disease symptoms were scored as illustrated: 1, no symptoms; 2, limited browning; 3, necrosis; 4, necrosis and gumming; 5, necrosis, gumming and spread of lesions from the site of inoculation. This experiment was performed once.

Inoculations of whole trees were scored for symptom appearance at and around the cut inoculation site (Fig. 3b). Only the two effectorless strains (ΔEff and ΔEffΔT) caused fewer symptoms at the inoculation site than the wildtype. Deletion of CEL effectors and Toxins (ΔCELΔT), or toxins alone (ΔT) greatly reduced the numbers of sites with dark necrotic lesions, gumming and spreading, but the effects were not statistically significant (Fisher’s exact test, p>0.05).

### Effectors and toxins work synergistically to promote disease in immature fruit

Stab inoculation of immature fruits showed that, unlike in other tissues, deletion of all effectors did not result in a failure to cause symptoms, unless the *syrsyp* toxin cluster was also deleted (Fig. 4). The effectorless mutant (ΔEff) recorded lesion diameters that were significantly reduced compared with wildtype, but only after 6 days of incubation. Deletion of the toxin clusters (ΔT) caused the same reduction in pathogenicity as seen with the effectorless mutant (ΔEff), highlighting the greater role of toxins in fruit symptoms. The ΔFΔP mutations did not reduce lesion diameters but the ΔFΔPΔC deletions together had a significant effect in reducing disease symptoms. As observed in experiments on cut shoots, the CEL and toxins deletion combination (ΔCELΔT) strongly reduced symptom formation, an effect attributed mainly to *syrsyp* since the deletion of the syringolin A genes (Δsa) only led to a small reduction in lesion size compared to the *syrsyp* (Δss) mutant, which could not form lesions.

**Figure 4.**
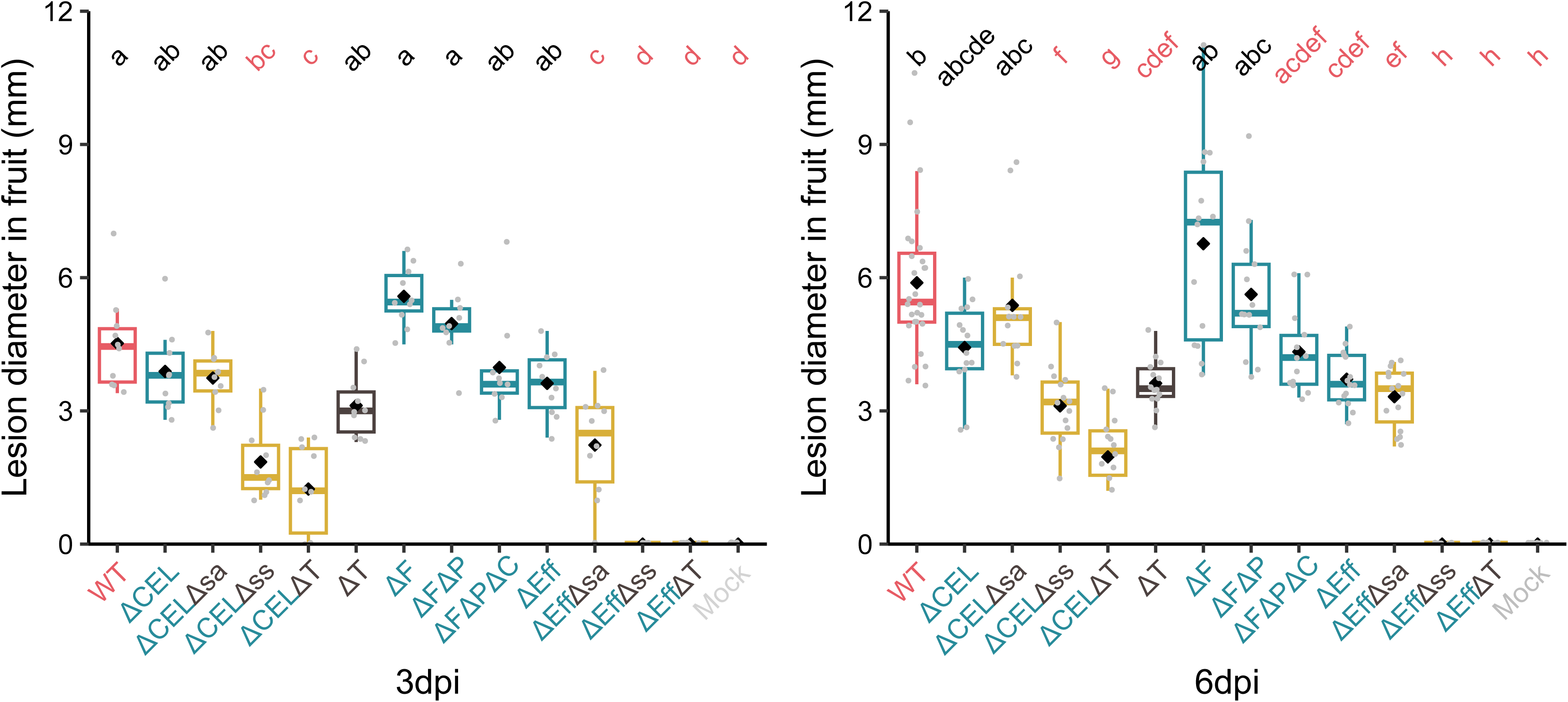
Lesion formation in immature cherry fruits of cv. Sweetheart stab-inoculated with wild-type *Pss*9644 and deletion mutants as described in detail in Table 1. WT: wild-type, CEL: Conserved Effector Locus, F: FLEXIBLE group, P: PRUNUS group, C: CORE group, Eff: all effectors in CEL, FLEXIBLE, PRUNUS, CORE groups, sa: syringolin A cluster, ss: syringomycin/syringopeptin cluster, T: both toxins clusters. Mock: sterile toothpick. Letters in common above data points indicate no significant difference between treatments. Letters in red indicate significant differences compared to wildtype (p<0.05). This experiment was performed twice and data from 10 fruits for each treatment were analysed after log transformation, with ANOVA and posthoc Tukey-Kramer HSD tests to assess pairwise differences between mutants. Representative symptoms are shown in Fig. S4.

### Effectors are key virulence factors to enable infection of cherry leaves

Two series of experiments were completed on leaves of sweet cherry cv. Sweetheart. The first focused on the strains with deletions of toxins, CEL and all effectors (Fig. 5a and 5b). The second focused on deletion of the intermediate effector groups CORE, PRUNUS and FLEXIBLE of the polymutant (Fig. 5c, 5d). We observed that deletion of CEL alone (ΔCEL) greatly reduced symptoms 3 days after inoculation, but this effect was overcome to some extent after 6 days (Fig.5a). The ΔF, ΔFΔP, ΔFΔPΔC sequential deletions had an additive effect on the reduction of symptom formation (Fig. 5c). Deletion of both toxin clusters (ΔT) reduced symptoms to a similar extent as ΔCEL after 6 days and this was primarily attributed to deletion of *syrsyp.* The CEL and Toxins deletion (ΔCELΔT) caused a striking reduction in lesion formation. The most pronounced change in pathogenicity was observed using the effectorless mutant (ΔEff), which failed to produce symptoms in leaves even when both toxin clusters were present. Supporting this conclusion, no symptoms were produced by the effectorless and toxinless mutant (ΔEffΔT).

**Figure 5.**
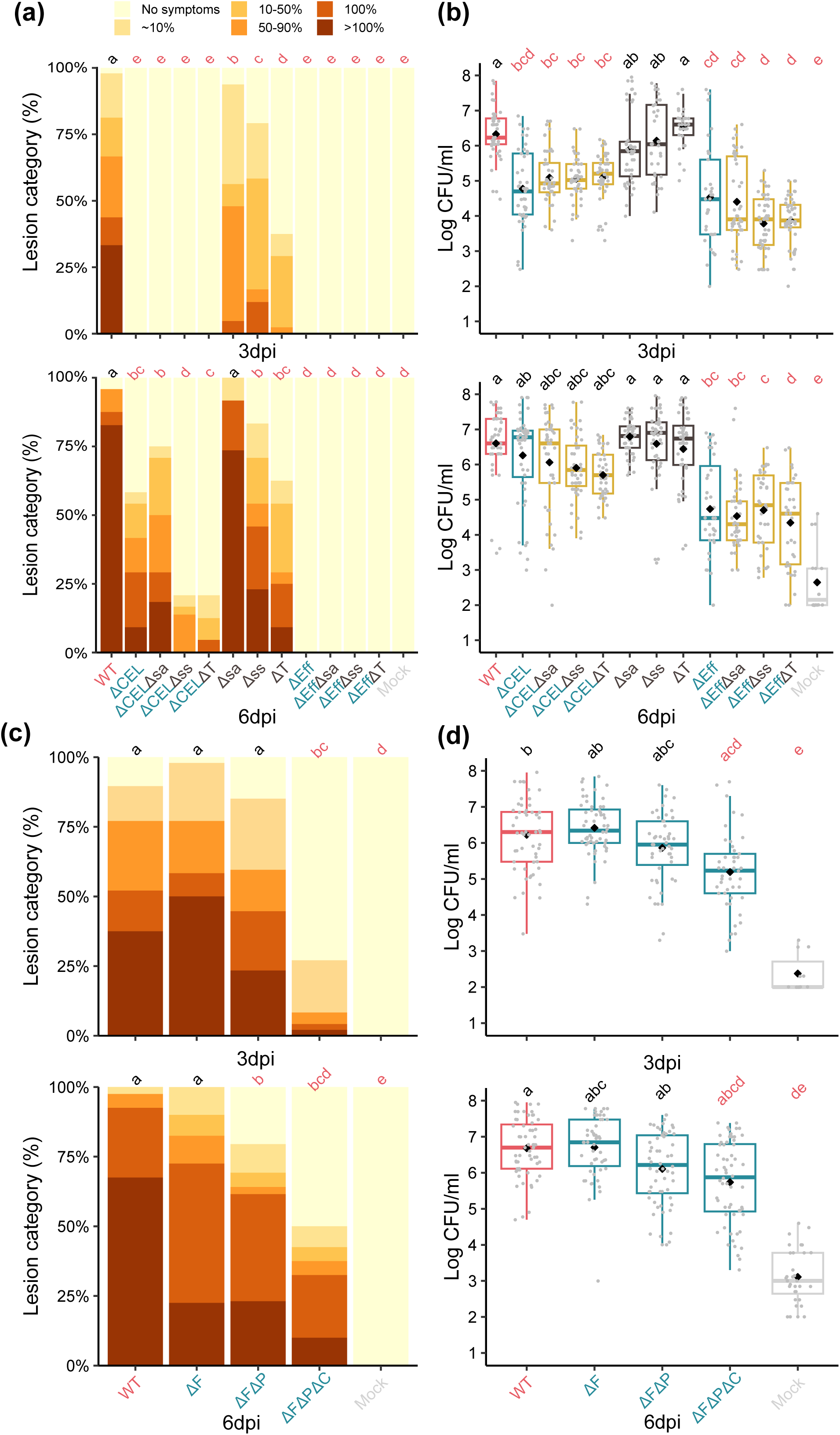
Comparison of the effects of mutations on the pathogenicity of *Pss*9644 to leaves of cv. Sweetheart and *Prunus incisa* using high concentration inoculum (2 x 10^8^ CFU per ml). Mutants are abbreviated as in Table 1, mock inoculation was with 10 mM MgCl_2_. Lesion formation was assessed as in Fig. 4Detached leaves of cv. Sweetheart infected with wildtype *Pss*9644 and deletion mutants as described in detail in Table 1, using low concentration inoculum (2 x 10^6^ CFU per ml). WT: wild-type, CEL: Conserved Effector Locus, F: FLEXIBLE group, P: PRUNUS group, C: CORE group, Eff: all effectors in CEL, FLEXIBLE, PRUNUS, CORE groups, sa: syringolin A cluster, ss: syringomycin/syringopeptin cluster, T: both toxins clusters. Mock:10 mM MgCl_2_. Letters in common above data points indicate no significant difference between treatments. Letters in red indicate significant differences compared to wildtype (p<0.05). (a and c). Lesion formation, assessed using a six-point scale as illustrated, based on the percentage browning /blackening at the inoculation site; 0, no reaction; 1, up to 10%; 2, 10-50%; 3, 50-90%; 4, 100% discoloration; 5, symptoms spreading from the infiltrated area. Data from three repeated experiments with 15 inoculation sites in total per treatment were analysed using pair-wise Fisher’s exact test. Representative symptoms are shown in Fig. S5. (b and d). Enumeration of *Ps* bacteria recovered from inoculation sites. Data are from three repeated experiments with 15 sites for each treatment per timepoint, were analysed after log transformation by ANOVA and posthoc Tukey-Kramer HSD tests to assess pair-wise differences between mutants.

An analysis of bacterial numbers at inoculation sites was carried out to gain a further insight into the impacts of deletions on bacterial fitness. Bacterial multiplication did not fully reflect the loss of symptom production observed (compare Figs 5a,c and 5b,d). For example, although there was a trend towards reduced multiplication by sequential deletion of ΔF, ΔFΔP and ΔFΔPΔC, the reduction was only significantly different from wildtype for the triple mutant (ΔFΔPΔC) 3 days after inoculation (p=0.05). Deletion of the CEL cluster (ΔCEL) reduced populations after 3 days, but not 6 days after inoculation. The combination of toxin and CEL deletions (ΔCELΔT) did not further reduce bacterial multiplication and the toxin deletions, although reducing symptoms significantly, did not alone result in the pathogen being unable to grow. The effectorless mutant (ΔEff) multiplied, but to a very low population density.

### Comparing symptoms in sweet cherry (*Prunus avium*) and *Prunus incisa* shows that ornamental cherry can be a model for the dissection of the roles of T3Es

Although *P. incisa* leaves are resistant to low inoculum concentrations of *Pss*9644 (2x10^6^ per ml), at high concentration (2x10^8^ per ml) infections, the symptoms that develop and the bacterial growth mirror those produced by low concentrations in sweet cherry leaves indicating a susceptible interaction (Hulin et al, 2022).)Fig. 6 shows the deletion of effectors caused much clearer reductions in symptoms in *P. incisa* than sweet cherry. For example, the ΔF deletion led to significantly reduced lesion formation 6 days after inoculation and further reductions in symptoms were observed with ΔFΔP. Delayed symptom development was again observed with the CEL deletion (ΔCEL). Deletion of toxins (ΔT) caused less effect on symptoms in *P. incisa* than in sweet cherry, but the CEL and toxins deletion mutant (ΔCELΔT) still produced very few symptoms.

**Figure 6.**
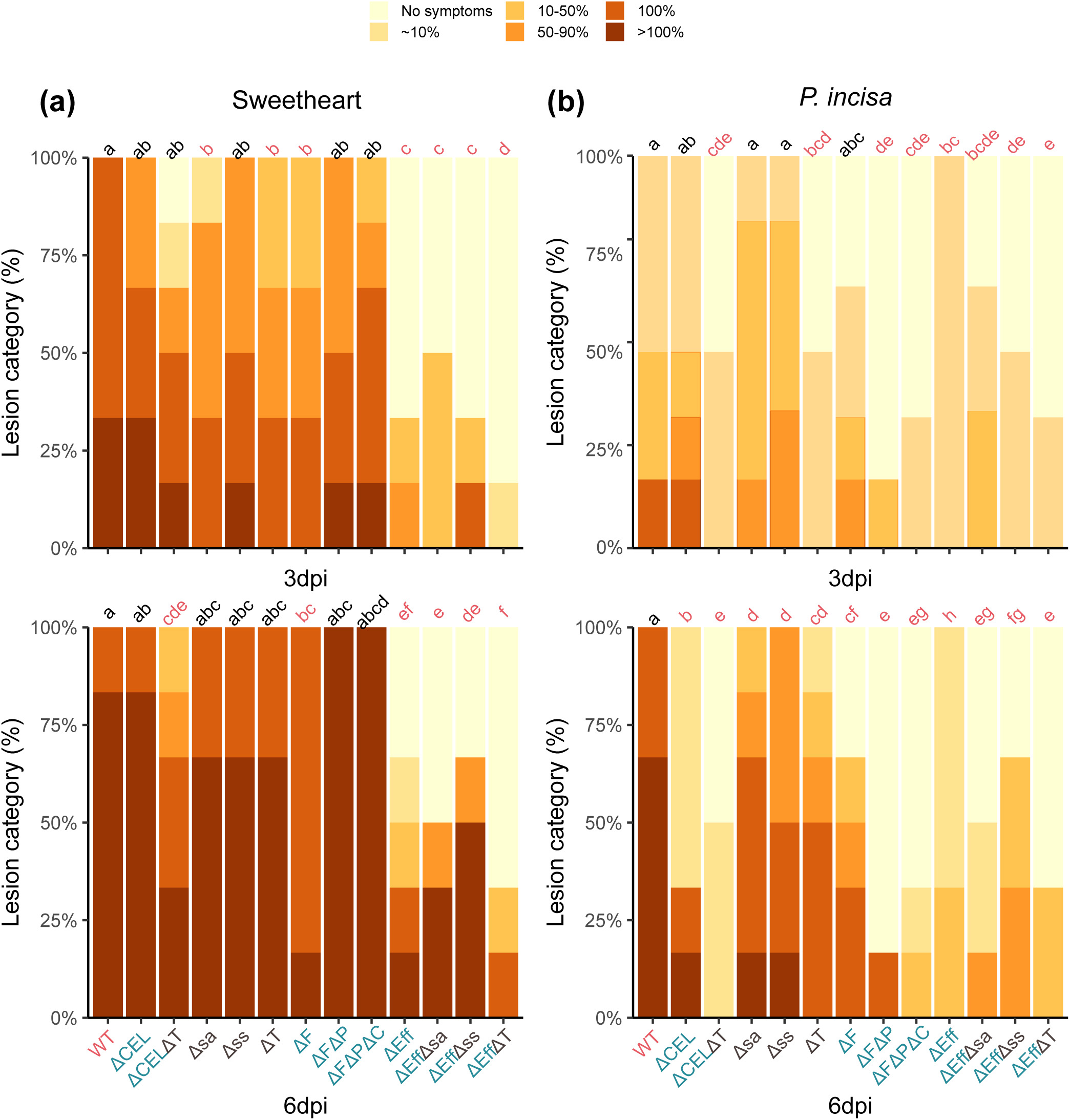
Comparison of the effects of mutations on the pathogenicity of *Pss*9644 to leaves of cv. Sweetheart and *Prunus incisa* using high concentration inoculum (2 x 10^8^ CFU per ml). WT: wild-type, CEL: Conserved Effector Locus, F: FLEXIBLE group, P: PRUNUS group, C: CORE group, Eff: all effectors in CEL, FLEXIBLE, PRUNUS, CORE groups, sa: syringolin A cluster, ss: syringomycin/syringopeptin cluster, T: both toxins clusters. Mock:10 mM MgCl_2_. Letters in common above data points indicate no significant difference between treatments. Letters in red indicate significant differences compared to wild-type (p<0.05). Lesion formation, assessed using a six-point scale as illustrated, based on the percentage browning /blackening at the inoculation site; 0, no reaction; 1, up to 10%; 2, 10-50%; 3, 50-90%; 4, 100% discoloration; 5, symptoms spreading from the infiltrated area. Data from three repeated experiments with 18 inoculation sites in total per treatment were analysed using pair-wise Fisher’s exact test. Note that at low inoculum concentration (2 x 10^6^ CFU per ml) *Pss*9644 fails to cause lesions in *Prunus incisa*.

### Effects of mixed inocula show that strains of *Pss* can co-operate *in planta* to cause maximal disease

A mixture of the effectorless mutant (ΔEff) that produces both toxins *syrsyp* and *sylA*, and the CEL-Toxins mutant (ΔCELΔT), which produces all effectors except those within the CEL, was examined to determine if the effect of the missing genes for toxin biosynthesis could be supplied, functionally, *in trans*. In theory, with complementation the mixture would have the same pathogenicity as the CEL mutant alone. Results presented in Figure 7 confirmed this hypothesis both in terms of symptom production (Fig. 7a) and bacterial multiplication (Fig. 7b).

**Figure 7.**
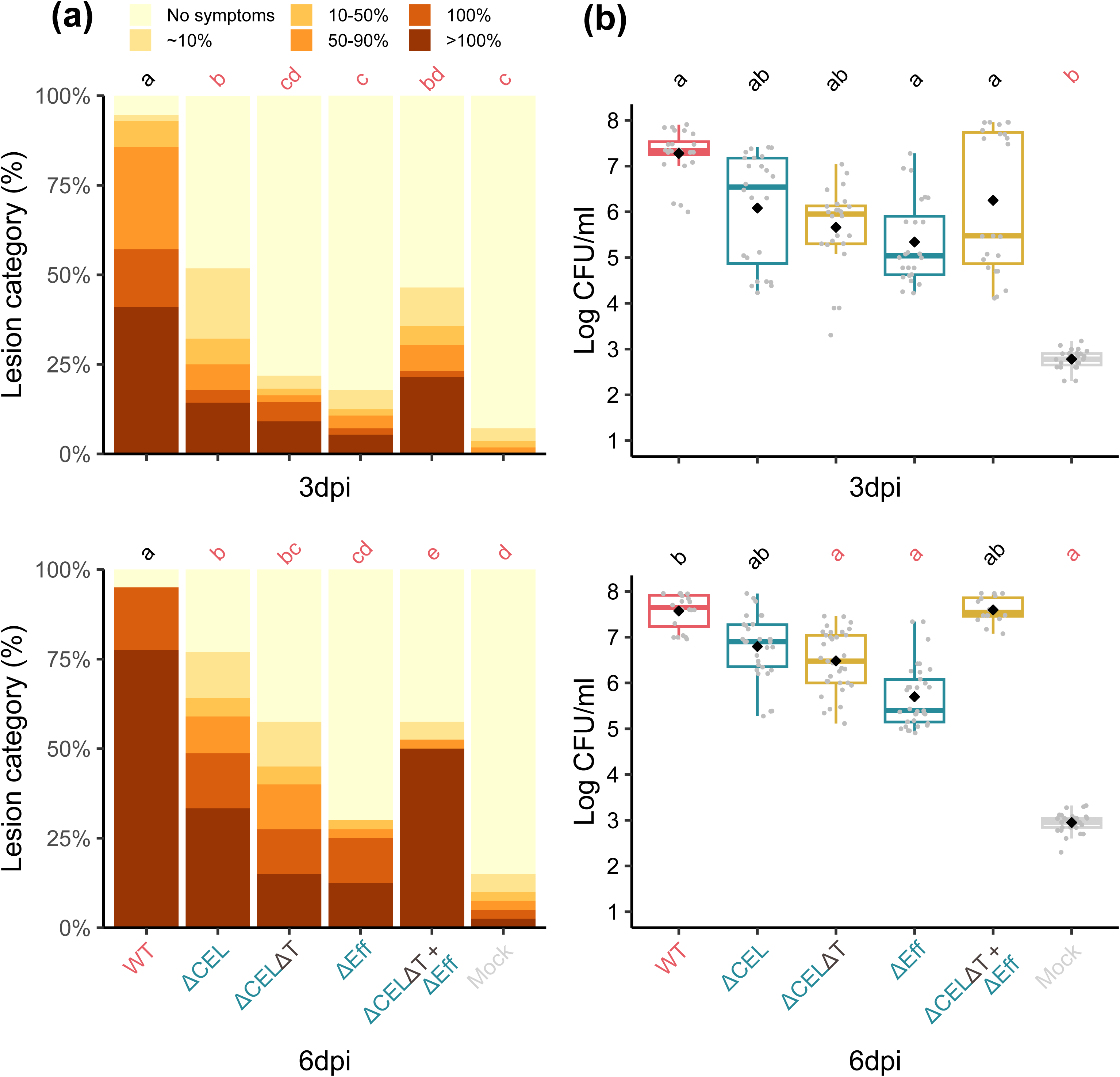
Use of mixtures of mutants of *Pss*9644 to demonstrate complementation of gene deletions. A mixture of the effectorless mutant (ΔEff) that produces both toxins, and the ΔCELΔT mutant which produces all effectors except the CEL was examined to complement the missing genes *in trans*. Pathogenicity to detached leaves of cv. Sweetheart was examined using low concentration inoculum (2 x 10^6^ per ml). WT: wild-type, CEL: Conserved Effector Locus, F: FLEXIBLE group, P: PRUNUS group, C: CORE group, Eff: all effectors in CEL, FLEXIBLE, PRUNUS, CORE groups, sa: syringolin A cluster, ss: syringomycin/syringopeptin cluster, T: both toxins clusters. Mock:10 mM MgCl_2_. Letters in common above data points indicate no significant difference between treatments. Letters in red indicate significant differences compared to wildtype (p<0.05). (a) Lesion formation, assessed using a six-point scale as illustrated, based on the percentage browning/blackening at the inoculation site; 0, no reaction; 1, up to 10%; 2, 10-50%; 3, 50-90%; 4, 100% discoloration; 5, symptoms spreading from the infiltrated area. Data from three repeated experiments with 15 inoculation sites per treatment were analysed using pair-wise Fisher’s exact test. (b) Recovery of bacteria from inoculation sites. Data from three repeated experiments with 15 sites in total for each treatment, were analysed after log transformation by ANOVA and posthoc Tukey-Kramer HSD tests to assess pair-wise differences between mutants.

## Discussion

### Testing bioinformatics-based predictions

Comparative genomics of the available genomes of strains of *Ps* including all phylogroups identified four effectors that were gained in pathogens of cherry: *hopAR1, hopBB1, hopBF1* and *hopH1* (Hulin et al., 2018a). Of these, the *Pss*9644 strain used here contained only *hopAR1* and *hopH1*, and these genes were part of the PRUNUS group identified in Figure 1. Mutation of the entire PRUNUS group did not cause a significant reduction in symptom formation in wood, or fruit of sweet cherry, indicating that their deletion did not impact on pathogenicity as predicted. Perhaps effectors remaining in the other effector groups allow bacteria to remain pathogenic through redundant roles in disease development. However, a minor effect was observed in leaves of sweet cherry and more clearly in *Prunus incisa* with the combined FLEXIBLE and PRUNUS group deletions (in ΔFΔP) causing further reduction than FLEXIBLE alone (ΔF). The clear results from *Prunus incisa* therefore support an important role for *hopAR1* and *hopH1* in the wider infection of *Prunus* species.

### Redundant effector groups and the importance of CEL

Our mutation strategy focused on the deletion of groups of effectors identified from genomic analysis of strains within phylogroup 2. The CEL group, physically linked to the *hrp* gene cluster that encodes the T3 secretion system, has been identified as an important group of effector genes in several strains of *Ps* (Badel et al., 2003, Munkvold et al., 2009). The other groups we selected based on phylogenetic analysis - PRUNUS, CORE and FLEXIBLE - were not identified as REGs as conceptualised for *Ps* pv. *tomato* DC3000 by (Kvitko et al., 2009). However, our results highlight that the REG concept is equally applicable to *Pss*/*Prunus* interactions. Effector deletions in DC3000 also caused stronger effects in tomato than in *N. benthamiana*, a finding similar to the results obtained comparing sweet cherry with *Prunus incisa* and indicating that the roles of effectors may vary in different host species (Kvitko et al., 2009).

The roles of the effectors encoded by *hopAA1*, *hopM1* and *avrE1* within the CEL in *Pss*9644 have been studied in detail in other pathosystems. In the *Pto*DC3000-tomato system studying bacterial speck, a *hopAA1-1* allele deletion mutant reduced chlorotic lesion symptoms (Munkvold et al., 2009); HopM1 was also implicated in lesion formation, but not enhancement of bacterial growth (Badel et al., 2003). In *Pto*23, AvrE1 functions in both roles (Lorang et al., 1994). In *A. thaliana*, the reduction in virulence of a CEL deletion mutant was associated with enhanced callose deposition adjacent to bacterial colonies, indicating that the CEL may act to suppress MTI (DebRoy et al., 2004). In *Pto*DC3000, AvrE1 and HopM1 have been shown to act as early time point bacterial growth promoters by creating an aqueous apoplastic environment (Wei and Collmer, 2018). Roussin-Leveillee et al., 2022 found that increased water-soaking was due to the effectors redundantly inducing stomatal closure by upregulating ABA pathways in the guard cells. Effects on stomata, which are much more common in leaves, may help to explain the greater reductions in pathogenicity observed caused by CEL deletion in cherry leaves rather than fruits and woody tissue. However, HopM1 has also been reported to have a role in suppression of the MTI-mediated oxidative burst in *A. thaliana* and *N. benthamiana* 24h post infiltration (Lozano-Duran et al., 2014, Wei and Collmer, 2018). In kiwifruit bacterial canker caused by *Ps* pv. *actinidae* (*Psa*), only *avrE1* is required for full virulence in leaves and together with *hopR1* (a related non-CEL effector) promotes bacterial fitness and necrosis (Jayaraman et al., 2020). Interestingly, *hopM1* and *hopAA1* in *Psa* do not have a role due to truncation and pseudogenization events, respectively.

### Toxins and CEL effectors operate synergistically

Syringomycin and syringopeptin are lipopeptide phytotoxins synthesised by a non-ribosomal mechanism of peptide biosynthesis encoded by the *syrsyp* cluster. They form pores in the plant cell membrane, disrupt ionic potential and cause cell death (Bender et al., 1999). Caponero et al., (1997) and Scholz-Schroeder et al., (2001) reported that mutations in the *syrsyp* cluster preventing their biosynthesis reduced the symptoms produced by *Pss* on immature cherry fruit by 30-70% compared to the wildtype strain. We confirmed the effects of *syrsyp* on fruit but found less significant reductions in virulence in other tissues. Syringolin A is a peptide derivative synthesised by a mixed non-ribosomal peptide/polyketide route. Although it has been implicated as a pathogenicity factor in several plants as a proteasome inhibitor in *Arabidopsis*, wheat and bean (Schellenberg et al., 2010, Misas-Villamil et al., 2013, Dudnik and Dudler, 2014), deletion of *sylA* had no clear effect on the pathogenicity of *Pss*9644 to cherry.

The influence of toxins was most apparent in fruits. Indeed, unlike in other tissues, the effectorless mutant still produced lesions in fruits, probably through toxin secretion. Fruit tissues may be more sensitive to the toxins and/or toxin synthesis may be enhanced by metabolites in fruit. This proposal is supported by the findings of Mo and Gross (1991) and Quigley and Gross (1994) who recorded enhanced production of syringomycin in media containing plant extracts such as arbutin and fructose. Despite all of the effector genes within *Pss*9644 being components of the HrpL regulon (Lam et al., 2014, Shao et al., 2021) we observed clear differences in T3E gene expression in *hrp-*inducing media). Such differences imply the potential for further regulatory control in the plant and indicate that the role of effectors may be modified depending on their expression under conditions within specific plant tissues. Shao et al., (2021) have identified several regulatory networks controlling T3E gene expression and their work highlights that regulation within plant tissues remains to be fully understood both in terms of nutrient availability and the location of bacteria within expanding colonies. The differential roles of toxins and effectors in the ability of *Ps* strains to colonise a wide range of ecological niches could be explored by further genetic dissection.

In all cherry tissues, the deletion of toxins and CEL together led to a strikingly synergistic reduction in pathogenicity. Understanding the cause of this effect may unravel mechanisms of defence targeted by the pathogen. Clearly, the importance of each of the components of the *Pss*9644 CEL interacting with *syrsyp* requires further dissection. Deletion of toxins reduced the development of lesions but did not reduce bacterial multiplication in cherry leaves. Similar differential effects on symptoms and populations have been reported for the toxin coronatine in *N. benthamiana* (Chakravarthy et al., 2018) and also in *Arabidopsis* following syringe infiltration (Brooks et al., 2005).

### Comparing sweet cherry and Prunus incisa

Our findings present an overview of the complex redundancy of T3Es operating in *Pss*9644. Apart from effectors within the CEL group, we have not been able to identify others with essential functions for pathogenicity to sweet cherry. Given the strong bioinformatics-led predictions of the positive role for effector groups in the evolution of pathogenicity to *Prunus,* it is perhaps surprising that clearer reductions in pathogenicity to sweet cherry were not found using the sequential deletion strategy. By contrast, in *Prunus incisa* several effectors emerged as having important roles. The nature of the resistance of *P. incisa* to low concentrations of inocula has not been explored. The ornamental cherry is resistant to all canker producing strains including *PsmR1* and *PsmR2* as well as *Pss* (Hulin et al., 2022). One explanation for the broad-spectrum resistance is that *P. incisa* has a strong MAMP response to *Ps.* In consequence, multiple effectors may be required to suppress MAMP-induced defences. The effectorless strain should allow further exploration of effector redundancy. It could also be used to unpick some of the components of resistance in wild cherry and related *Prunus* which may prove useful for more informed approaches to resistance breeding.

Certain sweet cherry varieties have been found to possess significant resistance to bacterial canker in the field, for example cv. Merton Glory, but resistance is not clearly apparent following artificial inoculation (Crosse and Garret, 1966, Hulin et al., 2022, Hulin et al., 2018b). The lingering concern in this work is therefore that the rapid lab- or greenhouse -based infection assays on sweet cherry create conditions that are very favourable to the pathogenic strains of *Ps*. In consequence the roles of groups of effectors may not be as apparent as in *P. incisa*. Direct infiltration of inocula does not allow effectors involved in the entry of bacteria into plant tissues to be assessed.

### Mixed inocula and effector guilds

The experiment with mixtures of mutants demonstrated complementation of the *syrsyp* and *sylA* deletions *in trans.* The use of mixed inocula containing several strains each expressing single effectors has been developed for *Pto*DC3000, carrying *in trans* complementation to a higher level (Ruiz-Bedoya et al., 2023). They used a metaclone containing a mixture of 36 co-isogenic strains in an effectorless background. Each co-isogenic strain was individually unfit, but the metaclone was collectively as virulent as wild type. This approach has led to the proposal that effector “guilds” exist in which effectors redundantly target the same host process (Bundalovic-Torma et al., 2022). A similar approach is now possible using the effectorless mutant of *Pss*9644. Given the synergism identified between the CEL and toxins in the infection of cherry it would now be helpful to consider the *Pss* toxins as components of the effectorome. The probability that diseases may be caused by mixtures of weakly virulent stains of *Ps* has been discussed (Hulin et al., 2023) who identified low virulence isolates in the field. The ecological significance of mixed inocula merits further investigation.

## Experimental procedures

### Bacterial culture

Strains of *Pss*9644 deletion mutants used in this study are listed in **Table 1** and plasmids in **Table S1**. The incubation conditions for *Pss*9644 and mutants were 28°C, and 180 rpm when cultured in liquid broth. *Escherichia coli* strains were incubated at 37°C, and 200rpm when in liquid. Agar or broth of Kings medium B (KB) (King et al., 1954) or Lysogeny Broth (LB) (Bertani, 1951) were used. Antibiotics and X-Gal used for screening were done using kanamycin (Km, 50 μg/ml), nitrofurantoin (Nif, 100 μg /ml) and X-Gal (40 μg /ml).

### Comparative genomics of phylogroup 2 strains of *Pss*

*Pss*9644 was grown in KB broth for DNA isolation using the cetyltrimethylammonium bromide (CTAB) method (William et al., 2012). Quality controls were performed using Nanodrop, Qubit and agarose gel electrophoresis. For long-read sequencing *Pss*9644 (SAMN17034057), a MinION (Oxford Nanopore, Oxford, UK) was used. Genome assembly, and annotation were performed as Hulin et al., (2018a). The genome was deposited in NCBI (assembly GCA_023277945, SAMN17034057).

Additional genomes (2686) belonging to the *Ps* species complex (taxonomic group ID 136849) were downloaded from NCBI on 21^st^ March 2023. FastANI v.1.33 (Jain et al., 2018) was used to calculate average nucleotide identity between pairwise genomes and R scripting was used to build a dendrogram of relatedness and group genomes into >90% identity groups (Hulin et al., 2023). 363 genomes belonging to the same group as *Pss*9644 were kept for further analysis and *Pph*1448A (a phylogroup 3) strain was kept as an out-group. Genomes of low quality (>5% contamination, <95% complete and N50 < 40,000 bp) were removed as in previous work (Hulin et al., 2022), leaving 324 genomes. Panaroo v1.3.2 (Tonkin-Hill et al., 2020) was utilised to generate a filtered core gene alignment of 3,943,305 nucleotides from 3825 core genes. A Maximum Likelihood phylogeny was built using IQ-TREE 2.0.4 (Minh et al., 2020) with model GTR+F+I+G4.

T3E genes were identified across the set of genomes using tBLASTn (BLAST+ v2.13.0) (Altschul et al., 1990). A database of 14,613 T3E proteins (Dillon et al., 2019) was utilised to query each genome. Putative hits were kept if they were over 50% ID and 50% query length. Bash scripting was utilised to obtain up to five non-overlapping hits for each effector family. The percentage of alleles for each family present across cherry pathogens was calculated manually. For visualisation purposes, only effector genes in *Pss*9644 are presented in the phylogeny.

To identify non-ribosomal peptide synthetase clusters in each genome the program antiSMASH 6.1.1 (Blin et al., 2021) was utilised. Bash-scripting was used to extract hits corresponding to the syrinomycin-syringopeptin cluster and syringolin A.

Results were plotted on the core genome phylogeny using R packages ggtree (Yu et al., 2016), ggtreeExtra (Xu et al., 2021) and phangorn (Schliep, 2011).

### Expression of genes encoding effectors and toxins in *Pss*9644

The expression of genes encoding potential virulence factors was compared after 5h growth in KB and Hrp-inducing minimal medium (HMM, Huynh et al., 1989) from overnight subculture. Media were inoculated with bacteria grown overnight in KB. Total RNA was isolated as previously described (Moreno-Perez et al., 2021) from three replicate cultures and sent to Novogene Co., Ltd. (Cambridge, UK) for cDNA synthesis, library preparation including rRNA removal and paired-end 150 bp sequencing performed on Illumina NovaSeq6000 platform obtaining 2 Gb raw data per sample. Adapter sequences and poor-quality reads were trimmed using fastq-mcf v1.04.807 (Aronesty, 2013) and quality checked using FastQC v0.11.9 (Andrews, 2010). rRNA decontamination was performed using BBDuk software v38.18. Transcript alignment and quantification were performed using Salmon v1.9.0 (Patro et al., 2017) with the long-read genome (GCF_023277945.1).

Differential gene expression analysis was performed using the DESeq2 package v1.40.2 (Love et al., 2014) in R version 4.2 (RCoreTeam, 2022). Before the analysis, a minimum read cutoff of 50 was imposed to prevent false-positive l2FC values. The DESeq2 contrast function was applied to the experimental (HMM) and control (KB) groups to provide an overall change in gene expression with an adjusted p-value of 0.05. Effectors and toxin synthesis genes were identified as in the comparative genomics analysis. RNA data were uploaded to NCBI (GSE255102).

### *Pss*9644 markerless deletion mutants

Markerless deletion mutants of *Pss*9644 were obtained as described (Kvitko et al., 2009). Briefly, flanking regions upstream and downstream of the genes of interest (∼400bp) were amplified using Phusion™ High-Fidelity DNA Polymerase (Thermo Fisher Scientific, UK) (Phusion PCR). The purified Phusion PCR products were amplified again by splice overlap PCR (SOE PCR) to join the two flanking regions. The purified SOE PCR product was double-restriction enzyme-digested and ligated into a pk18mob*sacB* vector previously double-digested with the corresponding enzymes and dephosphorylated with rSAP (NEB, UK). The constructs were transformed into *E. coli* DH5α cells plated on LB containing Km+XGal for blue/white selection, and positive transformants were confirmed using colony M13 PCR and Sanger sequencing (Azenta, UK).

T3E and toxin genes were individually deleted from the chromosome via a double homologous recombination process previously described (Hmelo et al., 2015, Neale et al., 2020), with modifications. Triparental mating was performed using *E. coli* DH5α cells as construct donor and with pRK2013 as a helper plasmid. Recipient *Pss* strains, donors, and helper were mixed in a 2 ml Eppendorf tube at a 2:1:1 volume (OD600 1.5: 0.8: 0.8). After centrifugation the pellet was carefully resuspended, plated on a KB agar plate, and incubated at 30°C for at least 24h. Transconjugants were selected on KB+Km+Nif plates. Colonies were streaked on LBA-no salt-15% (w/v) sucrose plates for sucrose counterselection. Merodiploid colonies were replica-plated on KBA and KB+ Km. Deletion mutants were identified by colony PCR.

### Pathogenicity tests

All pathogenicity tests were performed in 2022 on the susceptible sweet cherry cv. Sweetheart based on Hulin et al., (2018b). *Prunus incisa* was also used for high inoculum detached leaf assays. *In trans* complementation experiments were performed in 2023, under the same conditions.

### Woody tissue

#### Cut shoots

One year-old dormant shoots were collected at NIAB-EMR, UK in January. Fifteen shoots per bacterial treatment were dip-inoculated in 2x10^7^ CFU per ml suspension for each of the three independent experiments. Shoots were placed in a plastic box for the first two weeks and then randomised on rack trays filled with sterile distilled water. Incubation for 9 weeks was at 17°C, 16h Light(L):8h Dark(D) cycles. Lesion lengths (mm) were measured from the cut end of the bark-peeled shoot.

#### Whole tree

Wound inoculations were conducted in controlled growth rooms (20°C, 16hL:8hD cycles) in February, on 1 year old saplings. Inoculation sites were surface sterilised and 2cm of bark was sliced off. 50 µL of 2x10^7^ CFU per ml inoculum was pipetted over the exposed dormant wood and covered with parafilm and tape. 10 biological replicates per bacterial treatment were used performing six inoculations per tree and incubation lasted 9 weeks before scoring. Disease was scored as lesion category of the wound site after peeling it (no symptoms, limited browning, necrosis, necrosis + gummosis, necrosis + gummosis + spreading). The experiment was performed once.

#### Fruits

Immature fruits were stab-inoculated using toothpicks that had been touched onto 2-day old colonies grown on KB plates. An unused toothpick was used as a mock control. Five fruits were stabbed per bacterial treatment, with two inoculations per fruit for each treatment. Independent experiments were repeated twice. Lesion diameters were measured with a caliper (mm) 3 and 6-days post inoculation.

#### Leaves

Leaves were detached 1-1.5 week after emergence and inoculated with 2 × 10^6^ CFU per ml for symptoms and bacterial population counts in Sweetheart and 10^8^ CFU per ml for symptom comparisons in Sweetheart and *P. incisa*. Five leaves were used as biological replicates per bacterial treatment, with four infiltration sites per leaf. Three independent experiments were performed. Leaves were incubated at 22°C, 16h L:8hD cycles on sterilised trays with filter paper moistened with sterile distilled water and covered with a plastic bag to maintain humidity. Lesion scores were taken at infiltration sites 3 and 6-days after inoculation scoring: 0, no symptoms; 1, limited browning; 2, browning <50% of the inoculated site; 3, browning >50% of the inoculated site; 4, complete browning; and 5, spread from the site of infiltration.

Bacterial multiplication was examined at 3 and 6-days post inoculation at one infiltration site per leaf by excising a 1 cm diameter leaf disc after surface sterilisation. Discs were homogenised individually in Eppendorf tubes containing 1ml 10 mM MgCl_2_ and two stainless-steel ball bearings in a 2010 Geno/grinder, 1 cycle 30s at 1200rpm. From the homogenate, a 10-fold dilution series down to 10^−5^ was performed with sterile 10 mM MgCl_2_ and 10 μl aliquots plated on LB medium. After 2 days of incubation at 22°C, individual colonies were counted to calculate CFU per ml.

### Statistical analyses

R version 4.2.2 was used for experimental design, statistical analysis, and figure preparation. For statistical analysis, continuous variables such as “lesion length in mm” and “population counts” were tested as log data, to reduce skewedness, with ANOVA and posthoc Tukey-Kramer Honestly Significant Difference (HSD) test analysis to assess pairwise differences between mutants. “Diameter of lesion length” for fruit datasets were handled differently due to a zero-inflation problem. All non-zero mutants were formally tested for significant effects using a series of t-tests. Fisher’s exact test was performed on the symptom category classifications for leaves and whole tree with the null-hypothesis being that there is no difference between distribution of lesion type for pairs of strains. The p-values were then adjusted for multiple testing using the Benjamini-Hochberg procedure. Letters showing significant differences (P < 0.05) were obtained using the cList function from the rcompanion package. Box plots where shown indicate minimum, first quartile, median (line), mean (diamond) third quartile, and maximum values with bars indicating outliers.

## Acknowledgements

AV, MH, RWJ, JWM, RJH were funded by BBSRC grants BB/P006272/1 and the Bacterial Plant Diseases programme, BB/T010746/1. We thank Mojgan Rabiey and Nichola Hawkins for discussions and NIAB glasshouse staff for their role in plant maintenance. Support was received from students from the BSPP summer student programme and Global Training-Novia Salcedo. The authors declare no conflict of interest.

## Author contributions

The research was conceived by RJH, RWJ, JWM and MH with later input from AV. Experiments were designed and analysed by AV, MH, JWM, RJH and RWJ. In addition, SL assisted with cloning, JC assisted with RNAseq analysis, NG assisted with statistics, and ZZ and AG with pathogenicity testing. AV and JWM drafted the manuscript initially, with later input from all authors.

## DATA AVAILABILITY STATEMENT

The data that support the findings of this study are available from the corresponding author upon reasonable request.

## Supplementary materials

**Figure S1.** Principal Component Analysis (PCA) comparing gene expression of *Pss*9644 triplicate samples grown up in rich media (KB) as a control marked in red and minimal medium (HMM) as treatment represented in blue.

**Figure S2.** Heat map with differential gene expression log2 fold change ratio, comparing genes of *Pss*9644 triplicate samples grown up in rich media (KB) as a control and *hrp*-inducing minimal medium (HMM) as treatment. Red colours represent genes that are upregulated in HMM compared to KB. Blue colours represent genes that are downregulated in HMM compared to KB. Numbers in genes represent their ID in the annotated genome.

**Figure S3.** Representative illustration of a lesion score on cherry woody tissue pathogenicity test at 8wpi.

**Figure S4.** Representative illustration of pathogenicity test in immature cherry fruits at 3dpi and 6dpi.

**Figure S5.** Representative illustration of pathogenicity test in cherry leaves at 3dpi and 6dpi.

**Table S1.** Plasmid vectors and constructs used in this study for *Pss*9644 virulence factors deletions in the genome.

**Table S2.** Primers designed and used for this study for *Pss*9644 virulence gene deletion in the genome.

**Table S3.** PCR program details used for the different stages of construct cloning and putative mutant check.

**Table S4.** Transcript per Million (TPM) of *Pss*9644 triplicate samples grown up in rich media (KB) as a control and *hrp*-inducing minimal medium (HMM) as treatment.

