## Supplementary figures and images for "Genetic dissection of the tissue-specific roles of type III effectors and phytotoxins in the pathogenicity of *Pseudomonas syringae* pv. *syringae* to cherry"

### Figure S1

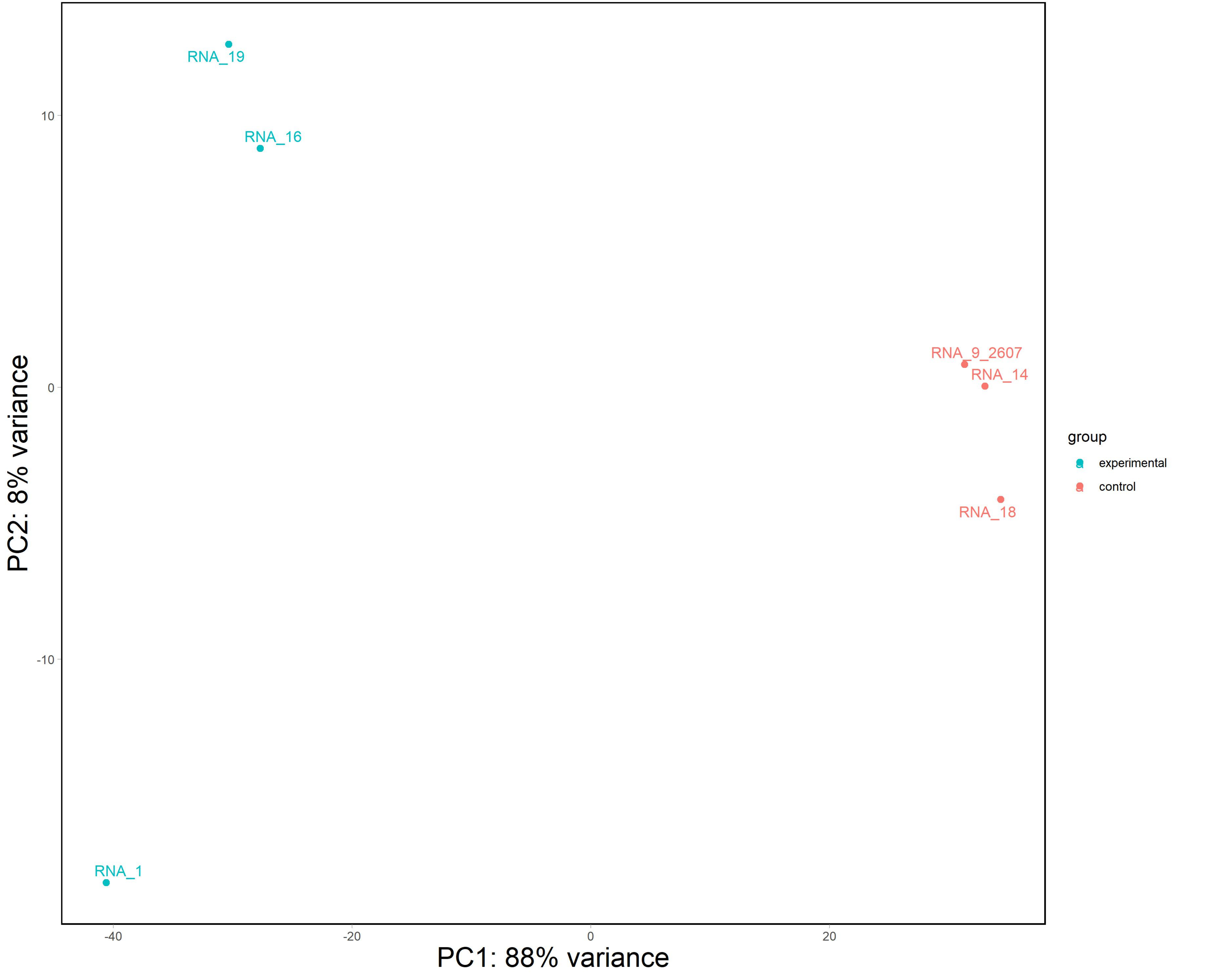

### Figure S2

#### Pss high l2FC

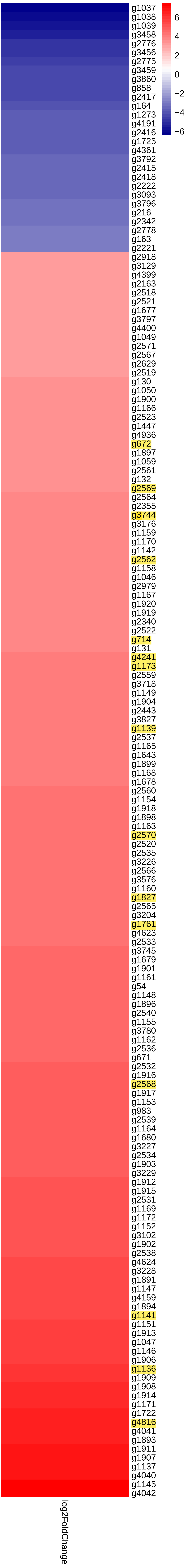

### Figure S3

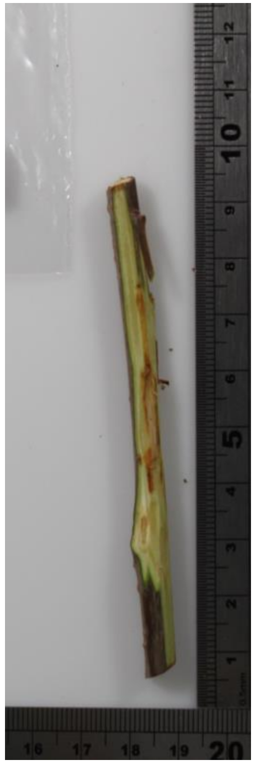

No symptoms

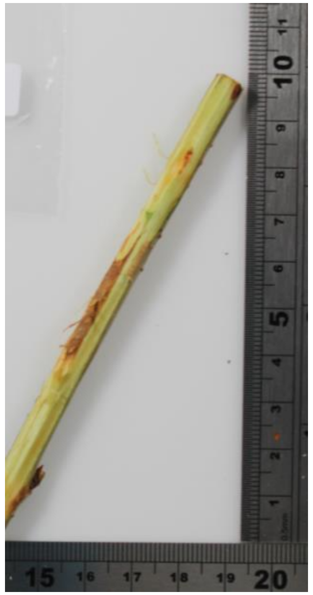

Browning

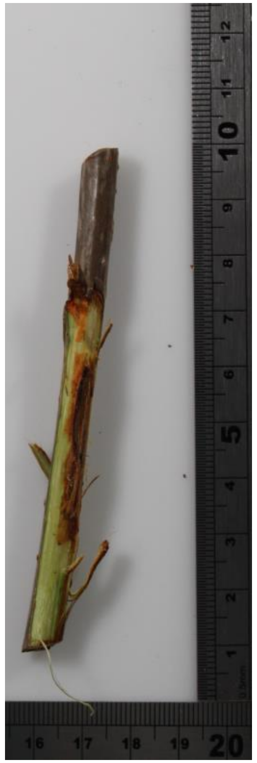

Necrosis

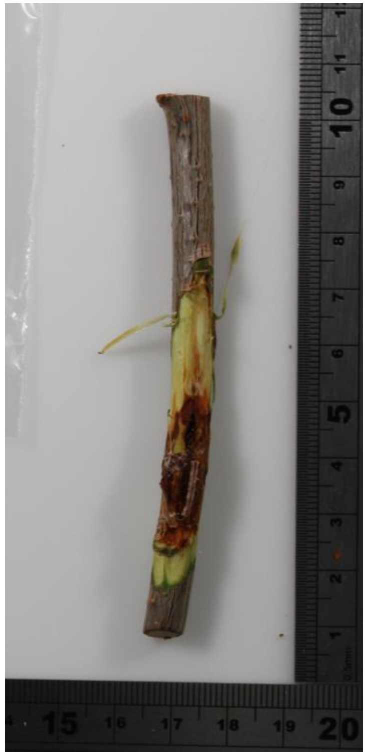

Necrosis and Gumming

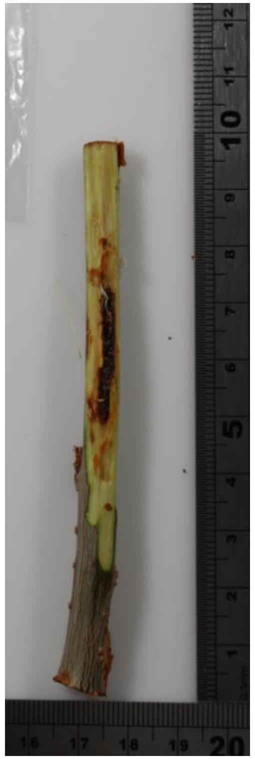

Necrosis, Gumming and spreading
