## Supplementary material for "Genetic dissection of the tissue-specific roles of type III effectors and phytotoxins in the pathogenicity of *Pseudomonas syringae* pv. *syringae* to cherry": Figure S4

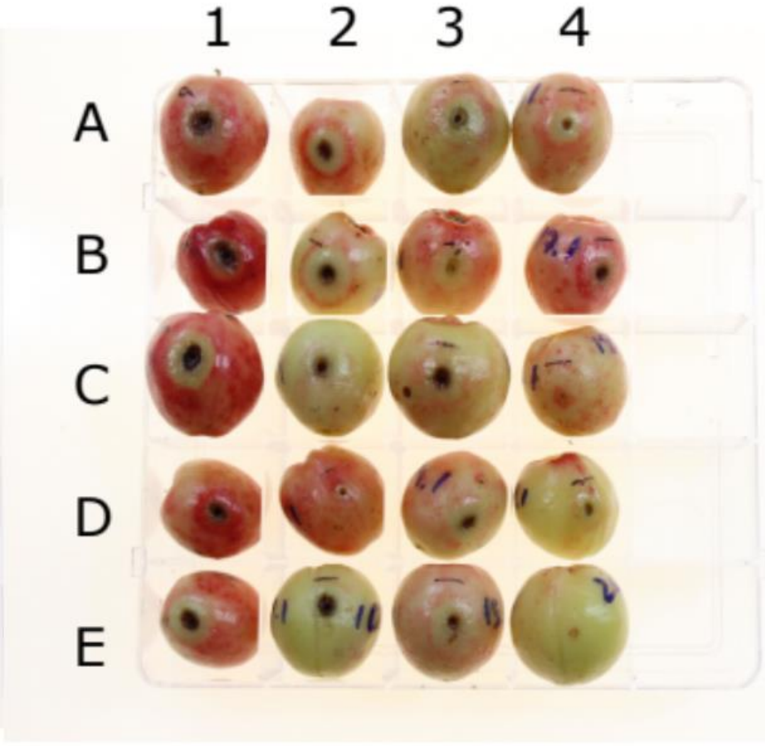


3dpi

1A: WT
1B: F
1C: FP
1D: FPC
1E: Eff
2A: Eff
2B: Eff
2C: hrpA
2D: EffT
2E: sa
3A: ss
3B: T
3C: CEL
3D: CELsa
3E: CELss
4A: CELT
4B: Effsa
4C: Effss
4D: Effsa
4E: EffT

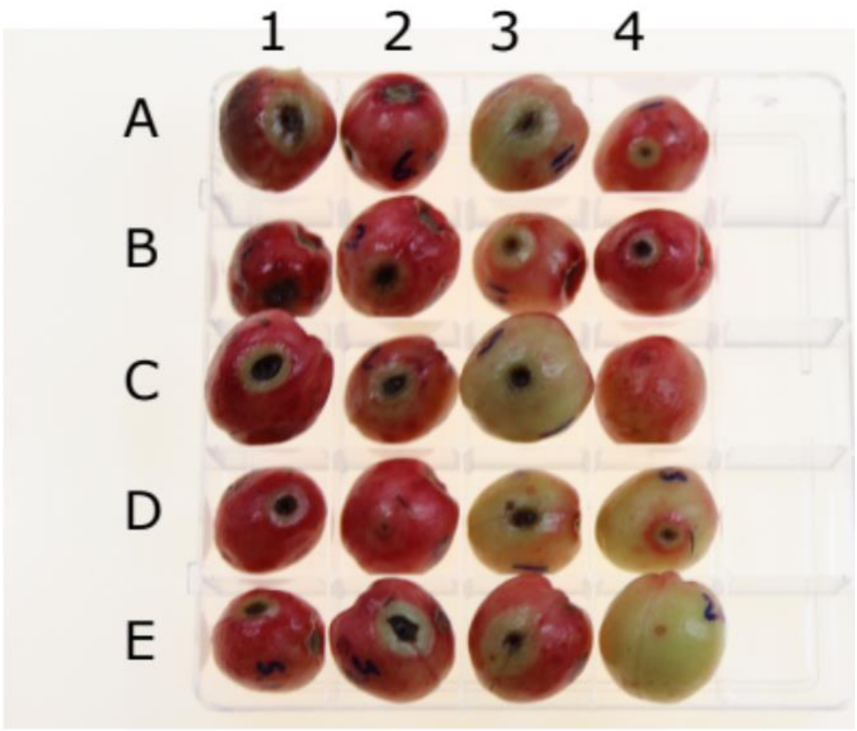


6dpi
